# A detailed graphical and computational model of the mammalian renal circadian clock

**DOI:** 10.1101/795906

**Authors:** Jessica R. Ivy, Barbara Shih, John B. Hogenesch, John J. Mullins, Tom C. Freeman

## Abstract

Here we describe the construction of a detailed graphical and computational model of the mammalian circadian clock. We use it to simulate the clock activity within the kidney, where it plays a pivotal role in regulating blood pressure. First, we assembled a network-based process diagram, which includes all known components of the clock and the interactions between them. Parameterisation of the model for Petri net-based simulation experiments used mRNA levels in the kidney to define initial conditions. With empirical testing, model parameterisation was further refined such that the simulated activity of core genes closely matched their measured activity. Furthermore, virtual knockout experiments performed on the model were shown to reflect experimental gene knockout data. It also identified points at which canonical clock genes may integrate with downstream genes likely to affect blood pressure and other aspects of kidney function. We believe that the model provides new insights into the complexity and function of this most central of physiological pathways and provides a valuable resource for the research community.

## Introduction

Endogenous circadian clocks are pervasive throughout the biosphere. They are an evolutionary adaptation to the Earth’s predictable daily cycle and associated fluctuations in light, temperature and associated activity patterns of an organism. As a molecular control system the circadian clock is active in every peripheral mammalian tissue (1, 2), orchestrating the cellular function according to the time of day. The molecular components at the core of the clock and the interactions between them appear to be highly conserved across all organ systems, but their downstream target genes are generally tissue- or cell-specific.

The renal circadian clock plays a pivotal role in regulating daily fluctuations in blood pressure (*3*). In healthy individuals, rhythmic variations in blood pressure are characterized by a ∼10% reduction during sleep followed by a surge in pressure upon awakening, with blood pressure plateauing during the day (*4*). Perturbations of this rhythm, particularly the nocturnal dip, confer increased risk for cardiovascular and renal disease (*5*). The kidney controls blood pressure, at least in part, through the modulation of sodium transport and extracellular fluid volume (*6*), which is brought about by specialized tubular epithelial cells lining the nephrons. These cells are heterogeneous, expressing different sodium transporters depending on their location along the nephron. In mouse the expression of many of these kidney transporters is altered following the knockout of the core clock genes, with alterations in the expression of the epithelial sodium channel (*Scnn1a* (*ENaC*)(*7, 8*)), sodium/chloride transporter (*Slc12a3* (*NCC*)(*7, 9*)), sodium hydrogen exchanger (*NHE3*)(*10*)), *Kcnj1* (*ROMK*)(*7*)), water channels (*Aqp2, Aqp3, V2R, V1aR* and *Aqp4* (*7, 11*)).

Genome-wide analysis of the kidney has established that a large number of genes expressed by the kidney exhibit a diurnal pattern of expression due the activity circadian clock (*2, 12*). Although the circadian rhythm of many of these urinary transporters has been well characterised, how these absorption/secretion pathways integrate with the renal clock is poorly understood. In order to better characterise the circadian-related biology of the kidney we decided to take a systems-based view - analysing its activity at the level of the transcriptome, modelling the interactions between core components of the mammalian clock and finally, simulating the pathway’s activity in this organ with and without perturbation.

Initially, we wished to understand the workings of the clock, but upon exploring existing models that describe this system, we found them to be overly simplistic (*13-15*). Indeed, no existing diagrams include all of its known components and the interactions between components are often poorly defined. Perhaps the best example available is present in the Reactome database (*16, 17*) (ID: R-HSA-400253), but this still lacks many details of the known biology of the system as described in the literature. There are also numerous mathematical models describing the circadian clock (*18-24*) and including a recent a kidney clock model (*25*). In design, these tend to be even simpler, at least in terms of their inclusion of the known molecular components and representation of the interactions between them, although the mathematics behind the models is often sophisticated.

Given the lack of a detailed model for this system, we set out to construct a pathway model of the mammalian circadian clock that reflects the consensus view of all known components of the system in man and the relationships between them. In order to achieve this aim, we undertook an extensive review of the literature, employing the mEPN modelling language (*26, 27*) to generate a graphical model of the clock, based on the principles of the process diagram (*28*). An advantage of the mEPN scheme over other graphical modelling languages is that it also directly supports the use of models for computational modelling, using a stochastic Petri net-based approach (*29*). In order that the model reflected the kidney clock, we then examined the CircaDB transcriptomics data (*2*) to provide us with a list of genes that are expressed in the kidney and identify those that are under circadian control. This information was then used to parameterise the model.

The outputs from this study include a detailed analysis of the circadian biology of the kidney transcriptome and a model of the peripheral mammalian circadian clock, which summarises the current knowledge of interactions between its components. Computational simulations using the model have shown it to closely recapitulate the activity profiles of genes expressed in the kidney and virtual knockout experiments of core components reflect experimental observations. The work also identifies the points at which canonical clock genes may integrate with downstream processes, regulating genes likely to affect blood pressure and other aspects of kidney function.

## Results

### Transcriptomic analysis

Gene expression data relating to the murine kidney were taken from CircaDB (*2*) and examined using a gene correlation network (GCN)-based approach. Clustering of the GCN identified a number of clusters of genes whose expression appeared to be under the regulation of the circadian clock, i.e. their expression coincided with the 12 h night/day cycle. When isolated from non-circadian regulated transcripts, the GCN was made up of two main groupings of genes. One set of genes being comprised of three main clusters (379 transcripts) whose expression was upregulated during the inactive period and another three main clusters (617 transcripts) whose expression was upregulated during the active period. The mouse being nocturnal, their active period is during the night. In both cases, clusters were comprised of genes whose expression peaked at different times during the day and night cycle. Of all the genes regulated, the largest relative change in expression over a 24-hour period was observed for many of the core clock genes, e.g. *Ciart, Bhlhe40, Nr1d2, Per1/2*, where expression peaked at the middle/end of the day (anticipating the active period). The expression of *Clock, Npas2* and *Nfil3* peaked during the middle of the night (anticipating the inactive period). The expression of the *Cry1/2* and *Rorc* peaked prior to end of the day and *Nr1d1* at the end of the night (Fig. 1). These expression profiles were used as benchmarks to optimise the parameterisation of the model. Gene Ontology (GO) enrichment analysis of the circadian-regulated gene clusters showed the greatest enrichment of any functional class of protein in the genes upregulated during the inactive period, were solute transporters and GO biological process enrichments also featured terms for transmembrane transport and circadian regulation of gene expression. In contrast those genes whose expression peaked during the night (active period) were enriched with genes whose GO function and process terms highlighted them to be associated with the response to unfolded protein and nucleoside binding (For full details of these analyses see Supp. Table 1).

**Figure 1:**
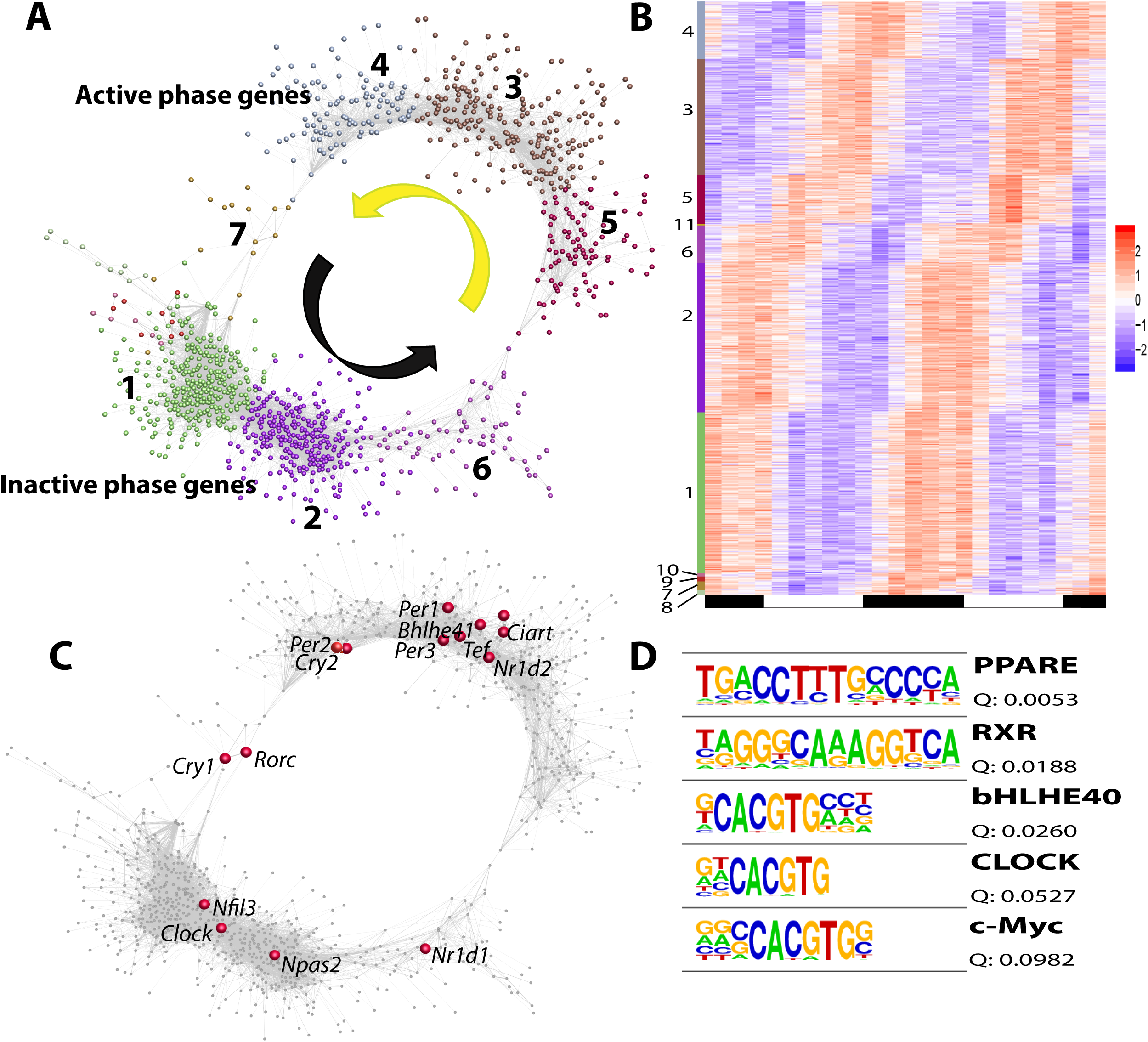
Network analysis of mouse kidney circadian clock driven gene expression. **(A)** Correlation network of 1061 transcripts (nodes) in the mouse kidney showing a diurnal pattern of expression. The structure of the graph divides the transcripts into two classes, those upregulated during the day (main clusters 3-5) in preparation for the murine active phase at night, and those upregulated at night (main clusters 1,2,6) in preparation for the murine inactive phase of the day. **(B)** Heat map showing the relative expression level (z-score) of the gene clusters shown in A. Dark/light bars at the bottom show the 12 h day/ night periods. **(C)** The GCN shown in A with the positions of the core clock genes shown. Note the sometime discrepant position in the network of gene paralogues, which in theory are regulated by the same factors. **(D)** Homer analysis of promoter enrichment in the active phase gene clusters. This confirms that many of the genes possess promoter elements for known clock proteins (bHLHE, Clock), but more surprising was the strong enrichment of PPAR promoters in these genes.

### Descriptive model of the circadian clock

A descriptive model of the circadian clock that includes all of the known components of the system and the interactions between them is presented in Figure 2. It incorporates 4 types of promoters, 21 genes, 51 proteins, 1 biochemical species and 43 protein complexes. Also included are 163 interactions, supported by 31 original papers and reviews (for a full list of these interactions and supporting evidence, see Supplementary Table 2).

**Figure 2:**
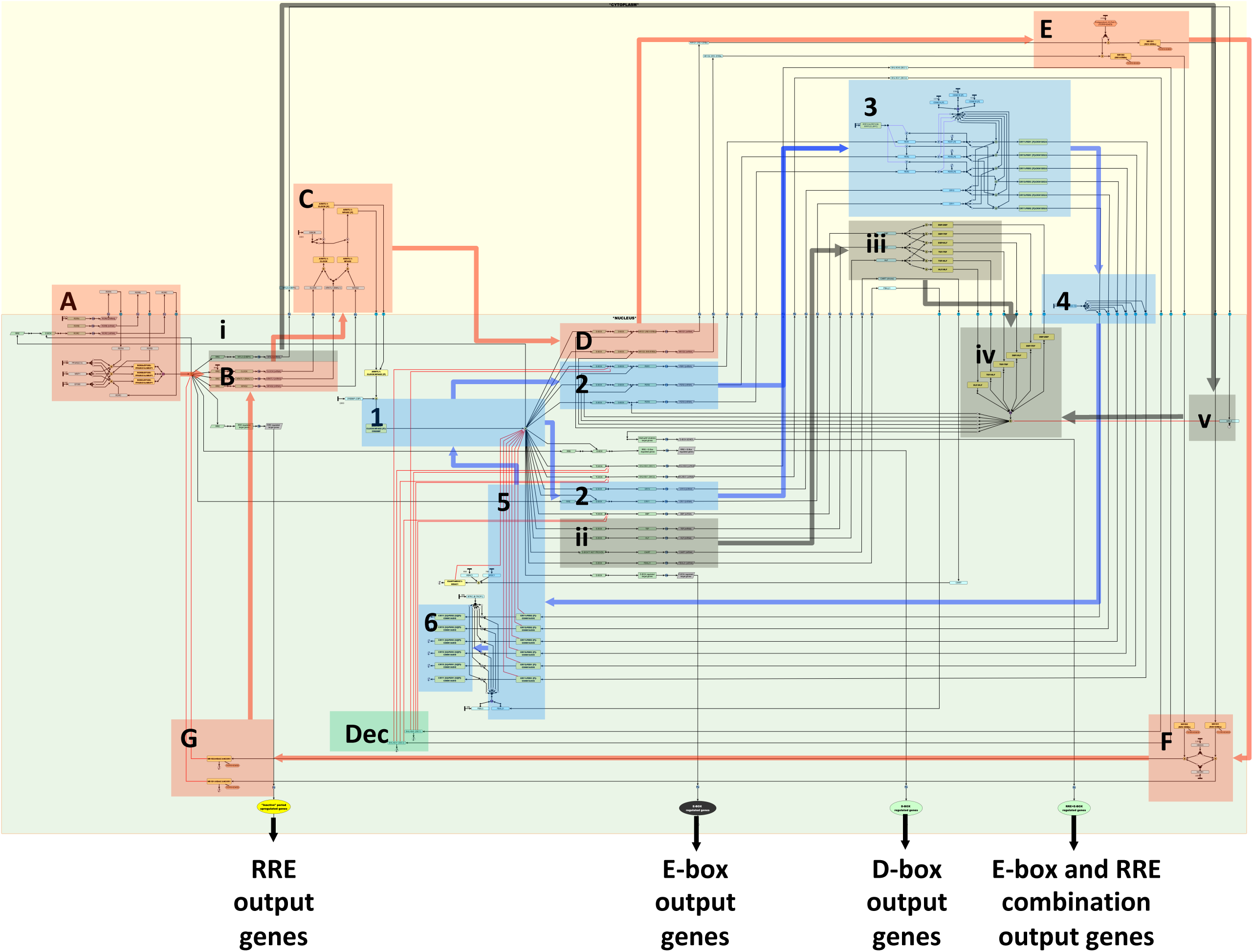
Descriptive pathway model of the mammalian renal circadian clock. Features of the classical ‘main’ transcription-translation feedback loop are highlighted by numbered blue boxes: **(1)** activation of transcription of *PERs/CRYs* by BMAL/CLOCK, **(2)** transcription and **(3)** post-translational modification of PERs and CRYs and formation of heterodimer complexes, **(4)** translocation of PER/CRY complexes to the nucleus, **(5)** inhibition of BMAL/CLOCK-induced activation of *PER/CRY* transcription and **(6)** degradation of PERs/CRYs. Main elements of the accessory loop are outlined in red boxes lettered A-E: **(A)** activation of transcription of *BMAL/CLOCK/NPAS2* by RORA and RORC, **(B)** transcription and **(C)** post-translational modification of BMAL/CLOCK/NPAS2; **(D)** activation of NR1Ds (rev-erb) transcription by BMAL/CLOCK/NPAS2 complexes, **(E)** NR1D1/2 binding to ferriheme, **(F)** translocation of NR1D1/2 back into the nucleus and formation of HDAC3, NCOR1, NR1D1/2 complexes, **(G)** inhibition of *BMAL/CLOCK/NPAS2* transcription. The third system associated with the pathway is demarcated by numbered black boxes and describe the **(i)** activation of NFIL3 by RORA or RORC activation of RRE binding sites, **(ii)** activation of PAR-bZIP genes by E-box binding of BMAL/CLOCK, **(iii)** dimerization of TEF, HLF, DBP **(iv)** translocation to the nucleus and activation of D-box genes by the dimers **(v)** and inhibition of D-box activation by NFIL3. Finally, the green box marked ‘Dec’ highlights the inhibition of ‘E-box promoters by BHLHE40/41 (DEC1/2). Transcriptional targets of the core clock genes are represented by the outputs shown at the bottom of the diagram. The yellow background represents the cytoplasm and green, the nucleus.

The clock descriptive model can be separated into two major interlocking transcription-translation feedback loops: the main or canonical loop and the accessory loop, outlined in blue and red, respectively (Figure 2). A third feedback loop (outlined in black) includes D-box regulators, which may act to reinforce the rhythms set by the main 2 loops. Activation or inhibition of clock genes is regulated by three types of clock-regulated gene regulatory elements (CREs); enhancer box (E-Box, CANNTG), D-Box enhancer (TTATG/(C/T)AA) and Reverb/ROR-binding elements (RRE elements). The combination of elements confers timing to gene expression, with E-box regulated genes activated at the start of the active period, RREs activated at the start of the inactive period and D-box regulated genes somewhere in the middle of the active-period (reviewed (*30*)). Many clock-regulated genes have multiple RREs, E-boxes and D-boxes, but the available information on these and how they operate is limited, so no attempt was made to include them in the model.

In the main loop (Figure 2, highlighted in blue), helix-loop-helix (HLHb)-PAS transcription factors NPAS2, CLOCK and ARNTL (also known as BMAL1 or MOP3) and ARNTL2 (BMAL2) form heterodimers. These proteins are phosphorylated in the cytoplasm by GSK3 (*31*). They translocate back into the nucleus where they bind CREBBP. The complexes then bind E-boxes, which activate the genes that control the negative arm of the main loop, which include *PER1-3* and *CRY1/2*. PER1-3 and CRY1-2 mRNA and protein accumulates during the active phase in the cytoplasm (*32*). The PERs undergo additional regulation as they are phosphorylated by CSNK1E/D and subsequently degraded (*33*). The phosphatase complexes PPP1C1A-C are ubiquitous protein serine/threonine phosphatases acting in opposition to CSNK1E/D blocking the phosphorylation of PER1/2 (*34*). The PERs and CRYs eventually accumulate enough to form complexes with each other and CSNK1E/D/A which can then translocate back into the nucleus aided by KPNB1 (importin-beta) and go on to inhibit the binding of ARNTL-CLOCK complexes to E-boxes, thus switching off their own transcription. The CRYs within the complex are ubiquitinylated by FBXL3 (*35*) and FBXL21 (*36*) and PERs by BTRC, and degraded (*37*). FBXL21 is itself regulated by the main clock as it has an E-box in its promoter (*38*). As the PER/CRY complexes levels decrease, inhibition of ARNTL/CLOCK or ARNTL/NPAS2 to E-box binding is relieved and the cycle can start again.

In the accessory loop (Figure 2, highlighted in red), HLHb-PAS transcription factor binds E-box elements in the promoters of *NR1D1, NR1D2* and *RORA/C.* RORA/C go on to bind PPARGC1A, NRIP1 and EP300, the complexes of which bind RRE’s to promote transcription of *NFIL3, CLOCK, ARNTL/2* and *NPAS2*. Meanwhile, the proteins NR1D1/2 bind ferriheme-B in the cytoplasm and translocate back into the nucleus where they form complexes with HDAC3 and NCOR1 (*39*). These complexes can then inhibit the binding of RORA/C to RRE’s conferring rhythmicity to the HLHb-PAS genes (*NPAS2, CLOCK* and *ARNTL*).

The third component of the clock (Figure 2, highlighted in black) is made up of the PAR-bZip genes (*DBP, HLF, TEF*), which activate D-boxes and NFIL3 (or E4BP4), which competes for and inhibits D-box activation. These genes are often regarded as “circadian output genes” as their rhythm is set up by the core clock (PAR-bZip genes are activated by E-boxes and NFIL by RREs) and act as transcription factors to impose circadian rhythm on any downstream genes with D-box elements in their promoters. There is also evidence that they feedback to the core clock, as several of the classical clock genes have D-boxes (including *PER1-3, NR1D1/2*). The cyclical behaviour of these genes likely feeds back to reinforce clock rhythms rather than modifying them.

Other recently discovered circadian genes have also been included in the model, i.e. *BHLHE40* (DEC1), *BHLHE41* (DEC2) (*40*) and *CIART* (Chrono) (*41*). The two *BHLHE* genes are under E-box regulation and are activated by ARNTL-CLOCK/NPAS2 complexes (*40*). Little is known about any post-translational modifications or additional protein binding, but these proteins are known to translocate to the nucleus where they have been reported to inhibit the ARNTL:CLOCK or ARNTL:NPAS2-induced activation of several E-Box genes, which include themselves, *PER1*, and *DBP* (*40*). CIART is also activated by ARNTL, CLOCK, NPAS2, although the presence of an E-box in its promoter has not been confirmed. CIART translocates into the nucleus where it binds HDAC3 and the glucocorticoid receptor, NR3C1. This complex then inhibits E-box activation by ARNTL-CLOCK/NPAS2 (*42*). These pathways add an additional layer of complexity and control to the clock.

### Parameterisation and validation of the model

Parameterisation of the model included the addition of delay motifs and input nodes, with the assignment of tokens representing the amount/activity of given components. The final parameterised model contains the same number of molecular species as the descriptive but a significantly increased node and edge count, i.e. 2679 nodes and 2761 edges. Gene expression data from the kidney taken from the CircaDB database was compared to the token accumulation patterns of the mRNA nodes during simulation of the model (Figure 4). Simulations, resulted in a similar pattern of activity to the gene expression profiles of the mouse kidney. In a number of cases, the peaks of expression were slightly offset between that predicted and observed, suggesting extra levels of regulation not captured by the model.

**Figure 3:**
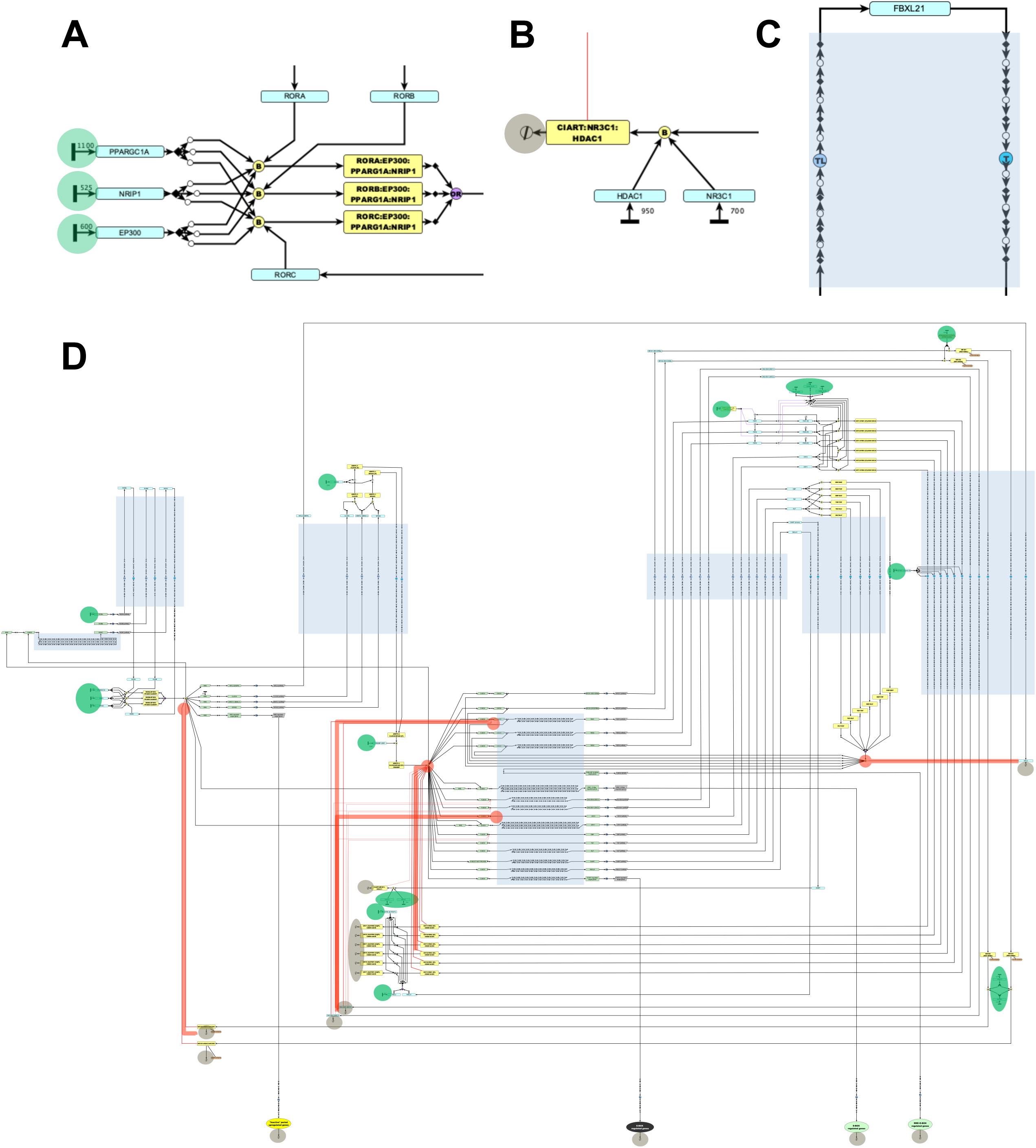
Parameterisation of the renal circadian clock model. **(A)** Token input points are highlighted in green and used where genes that are constitutively expressed in the kidney (as measured in the CIRCA kidney data). Token input levels at these points were based on their average expression level over a 24 h period and ensures that token levels at these points are maintained at a constant level throughout a simulation. **(B)** The inclusion of sink (transition) nodes and function as a means to lose tokens during a simulation. They represent the degradation of a protein and are indicated by red circles. **(C)** Delay motifs (repeat series of place and transition nodes) were added throughout the model, are indicated in blue. These represent multistep processes such as gene transcription, protein translation or translocation that are not defined in detail but take a finite amount of time. **(D)** All parameterisation points highlighted in the model.

**Figure 4:**
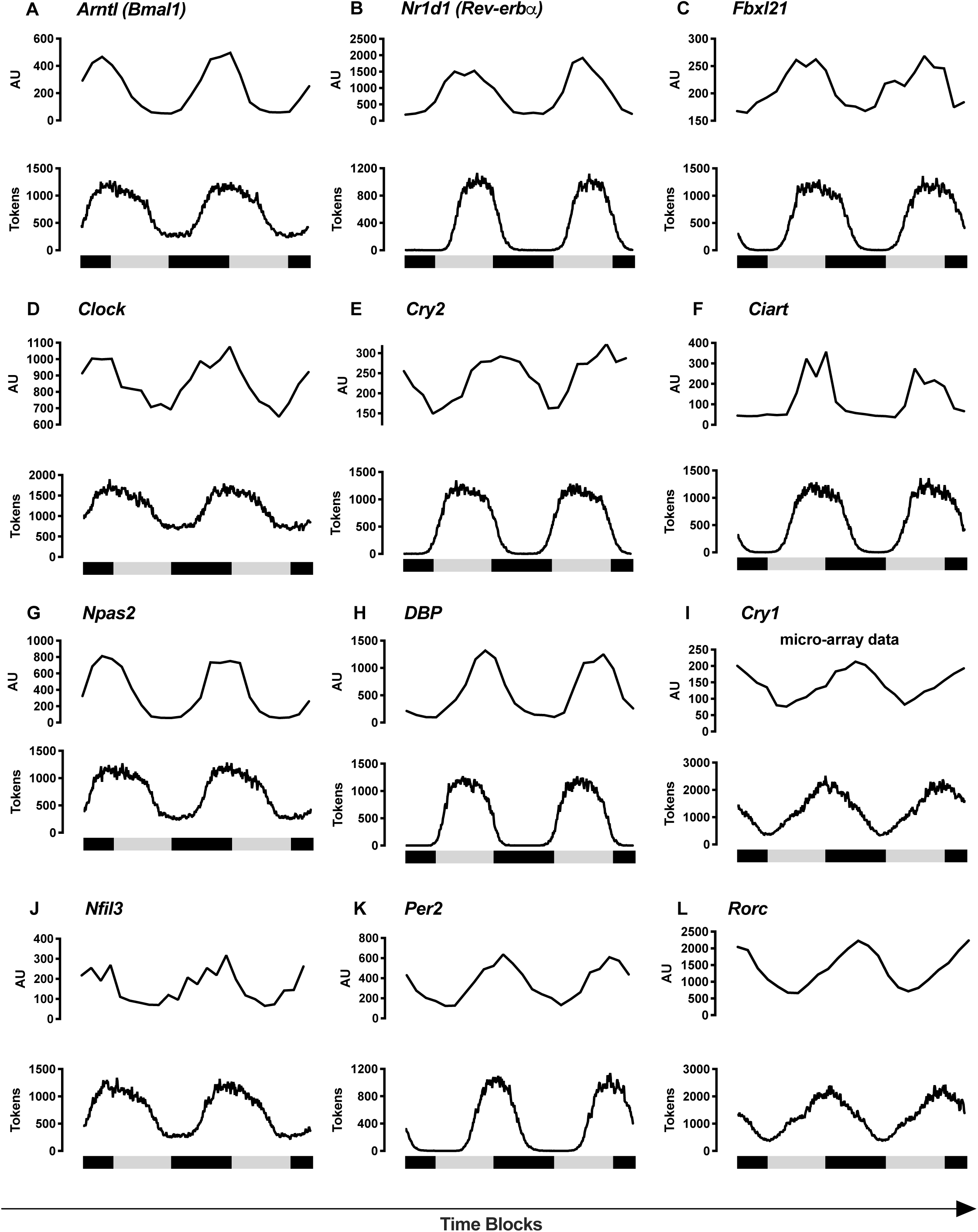
Comparison of the gene expression data and simulation data from the model of the mammalian renal circadian clock. Expression profile of core clock genes in the kidney (CIRCA-DB, top) compared to simulated gene activity profiles (bottom) as recorded during model simulation. **(A)** *Arntl*, **(B)** *Nr1d1*, **(C)** *Cry1*, **(D) (E)** *Rorc*, **(F)** *Clock*, **(F)** *Cry2*, **(F)** *Ciart*, **(J)** *Nfil3*, **(K)** *Nr1d1* and **(F)** *Rorc*. Below the plots, black bars represent dark/active period and grey the inactive/light phase.

### Knock out experiments

Several key genes were chosen to perform simulated knockout experiments. These were chosen based on the available peripheral knockout models in the literature and also on their importance in each of the main loops and accessory loops of the clock. As most clock genes have paralogues, we wanted to test the effect of knocking out a single versus both paralogues using the model. In our simulations, knockout of *NR1D2, NPAS2* or *CRY2* had subtle effects on the clock gene patterns (Figure 5A-C, grey lines). Double knockout of both genes from the accessory loop (*NR1D1* and *NR1D2*) resulted in arrhythmic, elevated *ARNTL* expression (Figure 5 Ai, red lines), but the oscillation of the tokens for *PER2* (Figure 5Aii, red lines) and *CRY1* (Figure 5Aiii, red lines) remained intact. Knockout of both functional paralogues from the positive arm of the main loop (*NPAS2* and *CLOCK*) blocked all other clock gene oscillations (Figure 5B, red lines). Finally, double knockout of both functional paralogues of CRY from the negative arm of the main loop (*CRY1* and *CRY2*) also suppressed all oscillations (Figure 5C, red lines).

**Figure 5:**
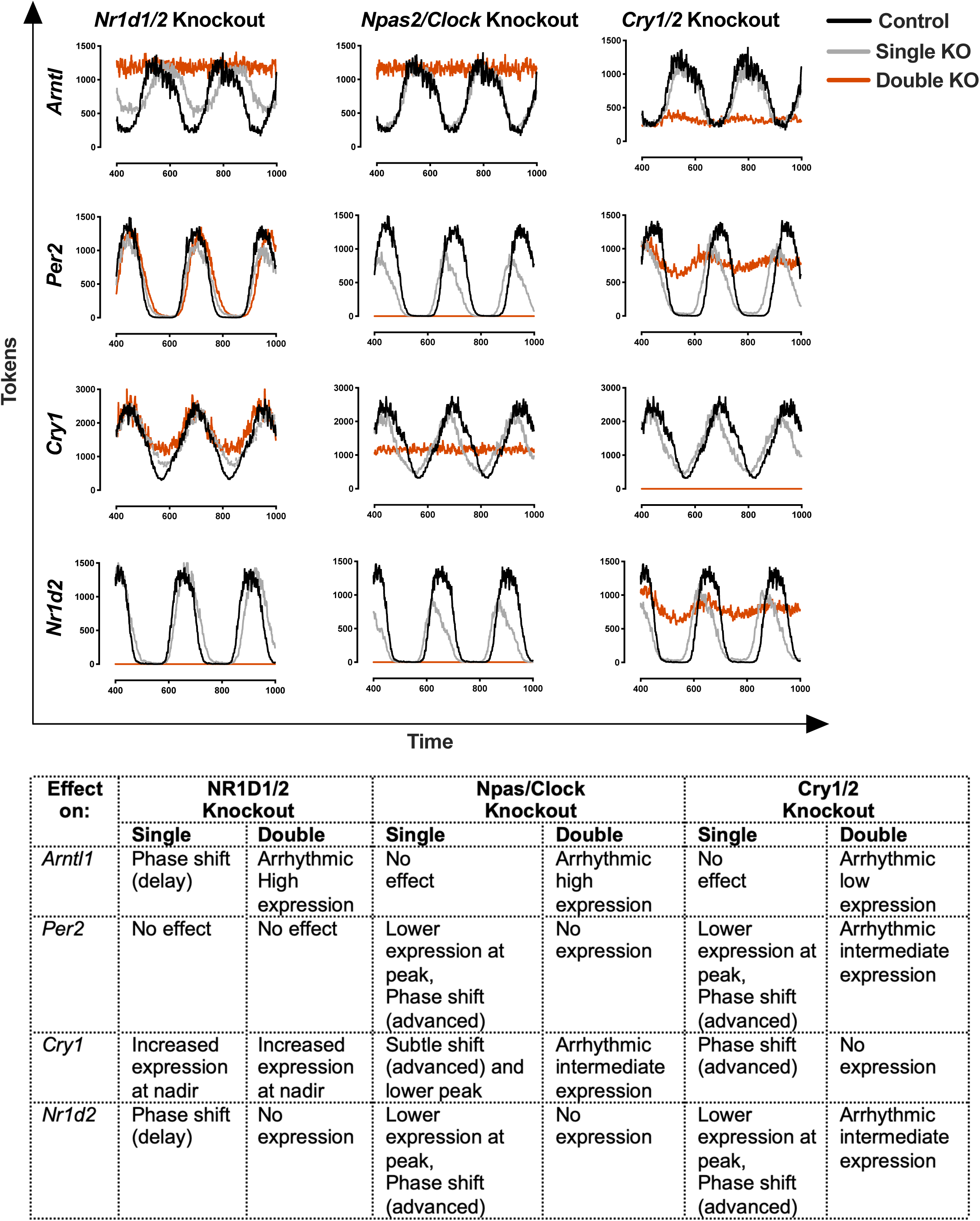
Results of Simulated Knockout experiments. Simulations of single and double knockout of circadian clock gene paralogues were performed using a computational model of a peripheral clock, parameterised for whole kidney. Gene/s knockouts are provided at the top of each graph set (column) and the effect of KOs on selected circadian clock genes are described on the y-axis (row). Results of the simulation experiments are described in the table below.

## Discussion

The kidney and the bodily functions it serves to regulate, exhibit a strong diurnal fluctuation in their activity. This is underpinned at the level of the transcriptome, which shows a high degree of circadian rhythmicity, our analysis showing 379 and 617 genes to be upregulated in during the inactive period (day) and the active period (night) of the mouse, respectively (Figure 1A and B). The level of rhythmicity in this organ is second only to the liver (*2*) and serves to tailor the functional activity of organs according to demand. In support of this, the greatest enrichment of any class of protein in the genes upregulated during the inactive period were solute transporters (n=34), their transcriptional activity increasing in preparation for the intake of food and drink, and the need to increase blood pressure during the active period (night-time in a mouse). Included in this list were *Aqp3* (water transporter), *Slc9a3* (sodium-hydrogen exchanger, Nhe3), *Slc2a2* (glucose transporter, Glut2). Strongly correlated with these genes were many known to encode core effector proteins of the clock machinery, e.g. *Bhlhe41, Cry2, Ciart, Per1-3, Nr1d2, Tef* (Figure 1C), and the promoters of these genes were highly enriched with transcription factor families known to be circadian regulators, i.e. Ppar, Bhlhe40, Rxr, Clock, Myc (Figure 1D). In contrast, the overarching GO enrichment of mouse kidney genes upregulated during the night, in preparation for the inactive daytime period, were associated with the response to unfolded protein and the metabolism of organic acids; perhaps indicating a more reparative activity of the organ during its ‘downtime’. This diurnal adjustment in the kidney transcriptome tunes its function to the different demands of it during the active and sleep periods. One of the fundamental roles of the mammalian kidney is to balance extra-cellular sodium in order to control and maintain blood pressure long term via a mechanism termed pressure ‘natriuresis’ (Reviewed: (*6*)). Importantly mutations in the core clock genes are accompanied by alterations in blood pressure (*8, 43*), (*44*), reviewed in (*45*)). Thus the inappropriate rhythmic regulation of the kidney is thought to contribute to the development of abnormal blood pressure rhythms. Therefore mechanistic insights into the workings of the autonomous kidney clock will be valuable in our search for better treatments to control blood pressure.

Despite its importance, relatively little is known about how the peripheral kidney clock operates. We therefore set out to build model of this system using a recently described graphical and computational modelling approach (*26, 27*). First, we examined the literature to define the major clock genes, referencing the CircaDB kidney expression dataset (above) to define their qualitative and quantitative expression in this organ. Guided by the literature as well as existing models (*13-17*), we then sought to define known interactions between components in order assemble a comprehensive model of the human peripheral circadian clock. In this model we aimed to include all known ‘core’ clock genes, including their lesser-characterised paralogues, e.g. *PER3* and *RORB/C* and molecules known to be specifically required in the post-translational modification of clock proteins or complexes of them. The use of the modified Edinburgh Pathway Notation (mEPN) scheme allowed us to describe graphically the interactions between all known clock components in unprecedented detail; the final model describing the interactions between 115 individual molecular species (promoters genes, proteins, complexes, biochemical) represented by 178 component nodes, and 163 interactions reported in the literature (Supp. Table 2) represented by 157 process nodes and 425 edges, 12 of which are inhibitory. The model includes three feedback loops; the so called classical ‘main’ transcription-translation feedback loop, the accessory loop and a third system that features the activation of TEF, HLF and DBP and their subsequent regulation of the REV-ERB and PER genes. Where appropriate, generic processes were explicitly depicted, e.g. the transcription, translation and translocation (albeit not in any detail), components were shown in the context of their sub-cellular location, i.e. nucleus or cytoplasm, and manual layout of the model was optimised for ‘readability’. All components included in the model were named using standard human nomenclature, although common names are also given, and component nodes were hyperlinked to the NCBI Gene database.

Whilst there have been numerous efforts to model the circadian clock previously (*13-17*), we believe this to be the most comprehensive model ever constructed of this system to date. Despite its complexity and our strenuous efforts to include all known players in this model, it is highly likely to be missing important elements. For example, little is known about the involvement of the peroxisome proliferator-activated receptors (PPAR) receptors in regulating circadian genes, yet circadian genes are highly enriched with promoter elements for them and *Ppard* shows a strong core gene-like regulation in the kidney.

Whilst the model described above is a highly valuable resource in its own right, in terms of providing a graphical overview of the current understanding of the mammalian circadian clock, given its complexity, it is impossible to know whether the system as depicted, could ever function. To test this, we sought to convert it into a computation model through a series of parameterisation steps (in reality the development of computation model went hand-in-hand with that of the graphical model, one informing the other). The mEPN language supports this through the introduction of nodes, motifs and tokens into a model that converts it into a form suitable for running Petri net-based simulations. When loaded into the network modelling software, Graphia Professional, it is possible to quickly run simulations using a stochastic Petri algorithm (SPN) (*46*) and visualise the activity of the system as a whole, or individual components of it. Through empirical testing of the model using an arbitrary number of tokens, we first tested the logic of the system, as inferred through the flow of tokens throughout all parts of the model. This involved ensuring the bipartite structure of the model was maintained throughout, i.e. alternating component nodes (places) and processes nodes (transitions), a perquisite for Petri net-based simulations. When this was done we observed that the system behaved as a damped oscillator. This is because not enough time elapses between activation and feedback inhibition for the system to reset. To compensate for this, delay motifs (linear series of alternating place and transition nodes) were added to account for multi-step processes such as transcription and translation, and served to stabilise the oscillatory behaviour of the model. Once the basic integrity of the model was confirmed, it was parameterised in line with the observed behaviour of its components in the kidney, as defined by the CircaDB transcriptomics data. Model components whose expression was essential for its integrity, e.g. in particular components whose presence was necessary for post-translational events and whose expression was for the most part stable, were given token numbers in line with their level of expression in the mouse kidney – gene expression being used as a proxy for protein levels. The only core gene where tokens were directly added to it was *Clock*, as a gene whose expression level showed some oscillatory behaviour but remained highly expressed even during its ‘off period’. We then set about trying to match the timing of a gene’s simulated activity to that observed in the data. To achieve this other delay motifs were added associated with a gene’s transcriptional machinery. For example, according to the literature the expression of *REV-ERB* and *PER* genes are regulated by the same factors, yet the timing of their expression is observed to be different and delay motifs were added to promoters of the *PER* genes to account for this. In these cases the delay motifs describe processes as yet unknown and in reality the differences observed in the timing of mRNA abundance, may not be associated with their transcriptional regulation as currently depicted.

Once fully parameterised, our model re-capitulated the behaviour of the peripheral clock, exhibiting oscillatory activity, with the simulated expression profile of many of its components matching that of the mouse kidney data. Moreover, the use of advanced visualisation methods to monitor token flow over time during a simulation makes understanding and appreciating some of the many complexities of the model far easier (See video – Supp. Data 3). We then wished to examine whether the model could predict the effect of knocking out core clock components in the periphery. Generating gene knockouts in the model is easily done by simply fixing the number of tokens to a gene to zero. A limiting factor in these comparisons is that there are currently only three kidney-specific clock knockout models described in the literature. The first is not directly comparable with our model as the *Arntl* (*Bmal1*) knockout is driven by the *Ren1d* promoter, which knocks *Bmal1* out only in the renin-expressing cells (*7*). This manipulation causes considerable effects on aldosterone and other endocrine signalling pathways that entrain and alter the intrinsic kidney clock, but does not inform us specifically about the intrinsic renal clock. The second, a *Per1* knockout, driven by the ks-cadherin promoter, does not detail the effects on the other clock genes (*47*). The third and most relevant for our purposes is an *Arntl* (*Bmal1*) knockout driven by an inducible *Pax8* promoter, which knocks out the gene in tubular cells (*48*). *Arntl2* (*Bmal2*) expression is very low in the kidney therefore the model does not include it. Our model predicts that the *Bmal1* knockout will behave in the same way as a double *Npas2/Clock* knock out, where the expression of all clock components will be arrhythmic: *Per2* and *Nr1d1* expression will be absent, *Cry1* expression intermediate and *Npas/Clock* levels will remain high. The data presented in the *Bmal1* kidney-specific knockout paper mirrored the model’s predictions with some exceptions. It displays low *Nr1d1* and *Per2* expression rather than their complete ablation, as predicted in our model. Pax8-driven *Bmal1* knockout is restricted to the tubule and so *Bmal1* is likely still expressed in glomeruli and blood vessels some *Nr1d1* and *Per2* expression here is to be expected and could account for the discrepancy. *Cry1* expression was arrhythmic and elevated, whereas the model predicted arrhythmic intermediate expression levels. The behaviour of *Clock* and *Npas2* was not described. Intriguingly, this *Bmal1* kidney knockout showed *Ppard* expression to be almost absent accompanied altered mitochondrial metabolism in kidney (*48*) reinforcing the observations made here with regards to its strong periodicity of expression and the enrichment of *Ppar* binding elements in the promoters of many kidney genes under circadian control. It is interesting to note that in liver also *Ppard* expression is also strongly periodic, as is *Ppara*, where their expression is strongly anti-correlated, suggesting a major role of members of this family of nuclear receptors in the regulation of the peripheral clock transcriptome of certain tissues.

Owing to the lack of other kidney-specific clock knockouts we made use of reports on other tissue-specific knockout models to test the computational model. *Nr1d1* (*Rev-erba*) knockout in the liver had no effect on *Per2* or *Cry2*, but elevated the basal levels of *Cry1* mRNA (as predicted by our model). Similarly, the liver knockout model showed elevated levels *Clock* expression, as observed also in simulations, we observed the nadir elevated with a small phase shift (*49*). The *Nr1d1/2* double knockout caused elevated baseline expression of *Cry1*, arrhythmic and elevated levels of *Arntl* and *Clock* mRNA, with a small decrease in amplitude in *Per2* expression but otherwise remaining rhythmic. All of these observations were mirrored by our simulation experiments. Single *Clock* knockout in explanted kidney slices resulted in rhythmic Per2 luciferase expression, but it was phase-shifted and had reduced amplitude (as predicted by our model) (*50*). Finally, in a double knockout of *Cry1/2, Per1/2* were observed to be elevated but arrhythmic, *Arntl* expression was very low and arrhythmic and *Nr1d1* exhibited intermediate expression (*43*). Again these observations were largely mirrored by the model, which showed arrhythmic *Per2* and *Nr1d1* expression with very low levels of *Arntl* expression. All in all, despite the fact that the model is highly complex in its structure, has been assembled based on information drawn from numerous studies in different species and tissues, often referring to the central clock (which at its core is thought to be similar), we have shown the model not only reproduces the normal behaviour of peripheral clock, but would appear to be highly accurate in predicting its behaviour following perturbation.

### Extensions and conclusion

In our search to better characterise and understand how kidney function and diseases associated with it and are regulated by the circadian clock, we have constructed what we believe to be the most comprehensive model of this system built to date. Furthermore, its presentation in a biologist-friendly manner, ease of assembly and computational analysis, means that it can be parameterised to represent other organs/cells and further complexity can be added and tested when experimental evidence becomes available. We have alluded to some of the likely missing or unknown elements of the system, for example, it is still unclear how having multiple and various combinations of RREs (*51, 52*) might affects kidney clock gene expression, but these could be added and tested. It is also well known that intrinsic peripheral clocks can be tuned by extrinsic cues through responses to entrainment factors, such as body temperature, glucocorticoids, and other hormones. The model currently includes the glucocorticoid receptor, NR3C1, where it binds with CIART and HDAC3 to inhibit the activation E-box containing genes. Glucocorticoids likely activate this receptor rhythmically and this could be one route that the steroids could tune the clock but further experiments are required to address how this may work and plasma glucocorticoid rhythms could be an important future input to add to the model. Furthermore, the model is primarily parameterised based on with mRNA levels but many renal transporters require post-translational modifications for activation and kinase, phosphatase and ubiquitinylase activity appear to also be rhythmic. It is currently unclear how these acquire their rhythms and indeed how or if they connect and integrate with the clock. These could be added to the model as and when the experimental data becomes available.

## Methods and Methods

### Transcriptome analysis of the circadian clock of mouse kidney

The CircaDB mouse tissue transcriptomics dataset (*2*) was downloaded from GEO (GSE54652) and data pertaining to the kidney isolated. A gene correlation network (GCN) analysis using the free network analysis software, Graphia Professional (Kajeka Ltd., Edinburgh, UK) was performed. Briefly, a .csv file of normalised expression values derived from the kidney collected every 2 h over a 48 h period were loaded into the tool. A correlation threshold of *r* = 0.85 was used to construct a GCN, where nodes represent transcripts and edges represent correlations above the set threshold. The network was clustered using the MCL clustering algorithm using an inflation value of 1.6 that defines the granularity of clustering. Clusters of genes exhibiting circadian regulation (a total of 1061 transcripts), were then reloaded into the Graphia and re-clustered (MCLi 1.6). An enrichment analysis performed on their associated Gene Ontology (GO) assignments and promoter elements using the tool ToppGene (*53*) on the clusters. For the full list of these genes, the kidney expression data associated with them and the results of the enrichment analyses see Supp. Table 1.

### Construction of a pathway model of the circadian clock

A network model of circadian clock pathway was assembled using information curated from Reactome pathway (ID: R-HSA-400253) and primary literature. The model was constructed using the network editing program, yEd (yWorks, Tubingen, Germany) and assembled according to the rules of the modified Edinburgh Pathway Notation (mEPN) scheme (*26, 27*). A full list of publications used in the construction of this model and the individual interactions they describe, can be found in Supp. Table 2. The core circadian clock components and many of the interactions between them have not been experimentally defined in the kidney, but are thought to be common to all peripheral tissues (reviewed in (*14*)). Whilst pathway components have been represented using human gene/protein symbols, we acknowledge that much of the known biology of this pathway has been defined in non-human systems. Therefore, this descriptive pathway model represents what may be considered to be a consensus view of the mammalian “peripheral clock” with evidence to support the model drawn from a range of systems.

The model was assembled in an iterative fashion. Initially, a simplistic negative feedback loop was constructed from the most basic components of the canonical clock, i.e. ARNTL, CLOCK, PER1 and CRY1. Through an extensive review of the literature, more complexity was added to the model, including proteins that do not oscillate but interact with or modify the clock proteins, e.g. PPARGC1A, NRIP1 and HDAC3. The “accessory loop” including NR1Ds and RORs was then added. Finally, paralogues were added including those of PER, CRY, NR1D, ROR. In some cases, there was no relevant literature defining the nature of the interaction of paralogues in the circadian clock. In these cases, the paralogues were ‘wired’ into the model to reflect their better-characterised counterparts. For example, PER3 regulation has not been extensively studied but it is thought to be important in peripheral tissues (*54*) and was integrated into the core clock in a similar way to PER1. ARNTL2, is thought to be a functional paralogue of ARNTL but is expressed at negligible levels in the kidney (*2*) and was thus excluded from the model.

Nodes representing individual proteins were hyperlinked to the relevant NCBI gene page via the URL field in the ‘Data Tab’ of yEd and similarly PubMed IDs relating to the evidence supporting a given interaction are included in the description field. All gene and protein components are labelled using their current human genome nomenclature committee (HGNC) symbol, but where a common name is widely used, e.g. ARNTL is commonly referred to as BMAL1 in the literature, this was also included, in round brackets () in a node’s label. Modifications to components such as phosphorylation and ubiquitinylation are included as additions to a component’s name (in square brackets), e.g. PER1 [P] for phosphorylated Period1 or PER1 [Ub] for ubiquitinylated Period1.

Expression of the core component genes is controlled by clock-regulated elements, present in their promoters (*55*). These include combinations of E-boxes, D-boxes and ROR-regulated elements (RREs) and were incorporated into the model. There are two different forms of E-boxes (*56*) namely E-boxes and E’-boxes but as the literature on these is sparse no attempt was made to distinguish between them in this model. BHLEH1 and 2 are reported to inhibit ARNTL/CLOCK activation of the E-boxes present in their promoters, as well as *PER1* and *DBP* and thus BHLEH1 and 2 were wired accordingly (*40*) otherwise all other E-boxes were treated in the same way. A full list of the literature sources used to construct the model and the interactions they define, are given in Supp. Table 2.

### Parameterisation

In order to parameterise the model ready for performing simulation experiments, a number of steps were necessary. Firstly, the pathway needed to conform to the specifications required of a Petri net, namely each node representing a molecular species (place node) is connected to a node representing a process or event (transition node) and visa versa, to ensure a bipartite graph structure. In cases where this was not the case, spacer place/transition nodes were added to the model. Secondly, ‘sink’ transitions were added to all nodes representing inhibitor proteins to simulate their decay (by proteasomal degradation), with tokens being lost from the inhibitor via the sink, e.g. the PER[U]:CRY[U]:CSNK1D complex is degraded in this manner, lifting the inhibition of BMAL1 complex binding to the E-Box genes. All inhibition edges in the model were of a competitive type (*26*). Thirdly, points at which tokens were required to be added to the pathway, i.e. proteins whose expression was not under the control of the clock itself, needed to be ‘marked up’ with tokens. Initially, this was done using an arbitrary number of tokens, to check that tokens were able to flow throughout the model i.e. to ensure there were no structural incongruences. However, once this was complete, token input was aligned with a component’s relative level of expression (as a proxy for the amount of the protein in the kidney), as defined in reference to the CircaDB gene expression atlas (*2*). For example, *Rora’*s expression did not oscillate and was expressed at around ∼625 normalised expression units, therefore 625 tokens were added as an input to the node representing this protein. In contrast, *Rorb* does not appear to be expressed in the mouse kidney, so it received no tokens in the model. *Rorc’s* expression does oscillate so its regulation within our model relies entirely on the flow input from the rest of the model.

Finally, delay motifs consisting of runs of place/transition nodes were added to the model. These represent multistep processes that are not defined in detail, i.e. transcription, translation and translocation or processes that are as yet unknown. The inclusion of delay motifs in the model, delays the time it takes for tokens to flow from one place to another. Without them the model exhibits damped oscillatory activity, that is to say the wave height on each round of oscillation decreases, as not enough tokens accumulate in each round to completely reset the system. Translation was given a consistent delay of a 6-place motif for all genes. Translocation of proteins/complexes back into the nucleus were given a minimum of a 14-place delay. Translocation was assumed to take longer in some instances, for example the translocation of CRY and PER complexes is thought to require accumulation prior to entry into the nucleus. It was also noted that activation of certain genes took slightly longer than others for reasons as yet unknown. For example, with reference to the CIRCA database, *Nr1d1/2* mRNA levels peak considerably earlier than *Per1-3* and *Cry2* mRNA despite in principle being activated by the same clock-regulated elements (E-BOX and D-BOX). *Cry1* and *Rorc* mRNA peak levels are even further delayed. This is supported by the literature: whereas *Per2* promoter and *Per2* mRNA have near identical activation profiles, *Cry1* expression is delayed by 4 h relative to its promoter (*57, 58*). We have added this delay to the model. *Rorc* and *Cry1* have near identical expression patterns and were therefore ‘wired up’ in exactly the same way. Parameterization of the model was also informed through extensive empirical testing and through further reference to the literature.

### Simulation of activity flow

Iterations of the model, as assembled in yEd, were saved as graphml files (the standard file format used by this tool) and loaded into Graphia Professional which incorporates the signalling Petri Net (SPN) algorithm (*46*). The tool’s graphml parser interprets the mEPN elements of the model, recognising component nodes as “places” and processes as “transitions” and converting the different mEPN nodes into their 3-D equivalents (*26*). Upon loading the model, the option is given to run the SPN algorithm to simulate activity flow. Simulations are made up of a series of time blocks during which all transitions are ‘fired’ once allowing tokens upstream of the transition to move to the node downstream of the transition being fired. A simulation is usually made up of number of runs, the results over a series of runs being averaged to give activity of a given component as tokens/time block. Enough time blocks are needed to move the tokens all the way through the pathway network. For example, ∼250 time blocks are required for this model to complete a full oscillation of the clock, i.e. one day. For all simulations 1000 time blocks and 200 runs were used as standard, with a uniform distribution and consumptive transitions.

Following a simulation, activity flow was visualized using the Graphia’s animation control tool. This represents flow through the nodes/places by changing both the colour (through a spectrum) and size (small to large) in line with the accumulation of tokens present over time. Activity flow was also visualized by use of Graphia’s built-in plot window providing a static graph of token flow over time in selected nodes (See video, Supp. Data 3). For more details on creating mEPN models and running simulations through them see (*26, 27*).

### Validation and knockout experiments

The validity of the model and its parameterisation was further tested through knockout experiments. These were performed by applying ‘0 tokens’ to the input edge of a gene target - this overrides any other input coming from other sources, with the effect that the gene is removed from the system. Run conditions and inputs were constant across each experiment and are detailed in Supp. Table 3. Results were compared to mRNA expression data from knockout experiments available in the literature. Few knockout experiments on the core clock genes have been performed in the kidney. Therefore, experiments performed in liver and other organ systems, were used as benchmarks against which the pattern of gene expression of the core clock genes following *in silico* knockout experiments was compared. Several papers were gathered for each knockout to generate a consensus on how the clock behaves under knockout conditions. Most mRNA species in the clock have paralogues, which function in a redundant manner, therefore we designed experiments to first knock out one gene, e.g. *Nr1d1-/-* and a second to knock out the paralogue as well, e.g. *Nr1d1-/-, Nr1d2-/-.* Six knockout experiments, including three single knockouts (*Nr1d1, Npas2, Cry1*) and three double knockouts (*Nr1d1/Nr1d2: Npas2/Clock, Cry1/Cry2*) were performed to test the major loops of the clock. The token input parameterisation for these experiments is shown in Supp. Table 3, all other simulation settings were as described above.

## Acknowledgements

JRI was funded by a British Heart Foundation (BHF) CoRE grant (BHF-RE/13/3/30183) and TCF is supported by a Roslin Institute Strategic Grant from the Biotechnology and Biological Sciences Research Council (BBSRC) [BBS/E/D/10002071].

## Author contributions

JRI and TCF constructed the models and performed the simulation experiments. BS helped with aspects of the transcriptomics analysis, and JBH and JJM advised on aspects of circadian biology and renal function. All authors assisted with the writing of the paper.

## Data Availability Section

The CircaDB dataset is available through GEO (GSE54652), all other information relevant to this work is included in the supplementary information.

