## Supplementary Information for "A detailed graphical and computational model of the mammalian renal circadian clock"

**Supp. Data 1. Descriptive model of the clock pathway**

An editable graphml file of the clock pathway descriptive model. To view and edit, download and open in the freely available network editing software, yEd (<https://www.yworks.com/products/yed>).

**Supp. Data 2. Parameterised model of circadian pathway**

An editable graphml file containing the parameterised computation clock pathway model. This can be opened and edited using the freely network editing software, yEd software (<https://www.yworks.com/products/yed>). Simulations can be run by opening the graphml file into Graphia Professional software (freely available at: <https://kajeka.com/download-graphia-pro/>). Once open you will be given the option to run a simulation, follow the interface to do this, e.g. using 1000 time blocks and 200 runs with a uniform distribution and consumptive transitions. Simulations can be visualised using the animation control function in Graphia Professional or simulation data can be opened for each node by selecting a node and using the shorthand control v. Alternatively, the simulation output data file can be downloaded visualised using a graph package. If node size is small upon loading, use shift + > to increase node size (or adjust using Node menu). More details on running simulations are available in (26, 27).

**Supp. Data 3. Video of circadian model flow simulation**

A video showing an example of a simulation animation from the fully parameterised circadian pathway model.

**Supp. Table 1. Circadian-regulated genes, mouse kidney**

Results of a gene correlation network (GCN) analysis. Included in this file are 4 tables (tabs): **1.** a non-redundant list day/night regulated genes; **2.** all mouse kidney data; gene enrichment analysis results for **3.** day and **4.** night upregulated genes.

**Supp. Table 2. Pathway Interaction Table**

A list of the individual interactions between molecular species presented in the descriptive model of the clock pathway, along with the supporting publications for each.

**Supp. Table 3. Pathway components and token addition**

A list of all the places where the pathway has been parameterised with token input. Tokens were only inputted for genes that were not rhythmic in kidney. Initially all inputs were run using 1000 tokens as the default input. This was then amended using the average values for these genes using CircaDB for kidney (2). Knockout experiments were performed by setting each test gene input as 0 this forces these genes to have their token level as 0. All other parameters were kept constant.
